## Supplementary figures and images for "Asymmetric Interactions Shape Survival During Population Range Expansions"

### paper_rw_0.1_Pwmm_0.5_Pmwm_0.75_no_num_psurv_multiple_rms.png

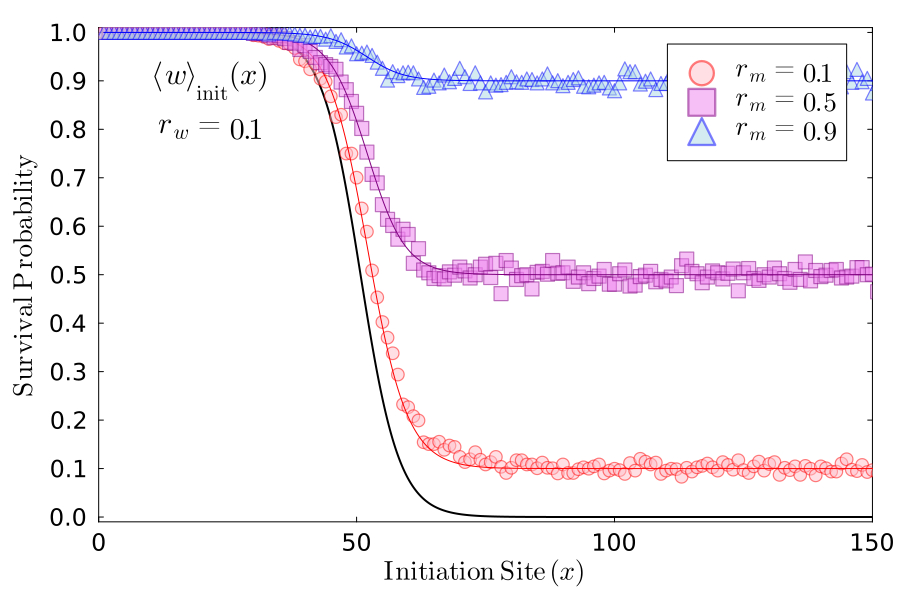

### paper_rw_0.1_Pwmm_0.5_Pmwm_-0.25_no_num_psurv_multiple_rms.png

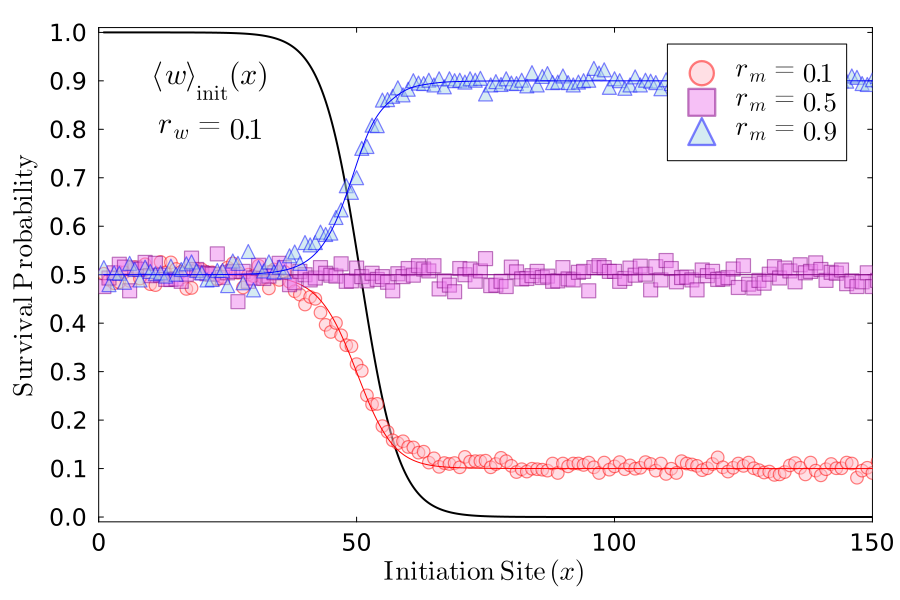

### paper_rw_0.1_Pwmm_-0.25_Pmwm_0.75_no_num_psurv_multiple_rms.png

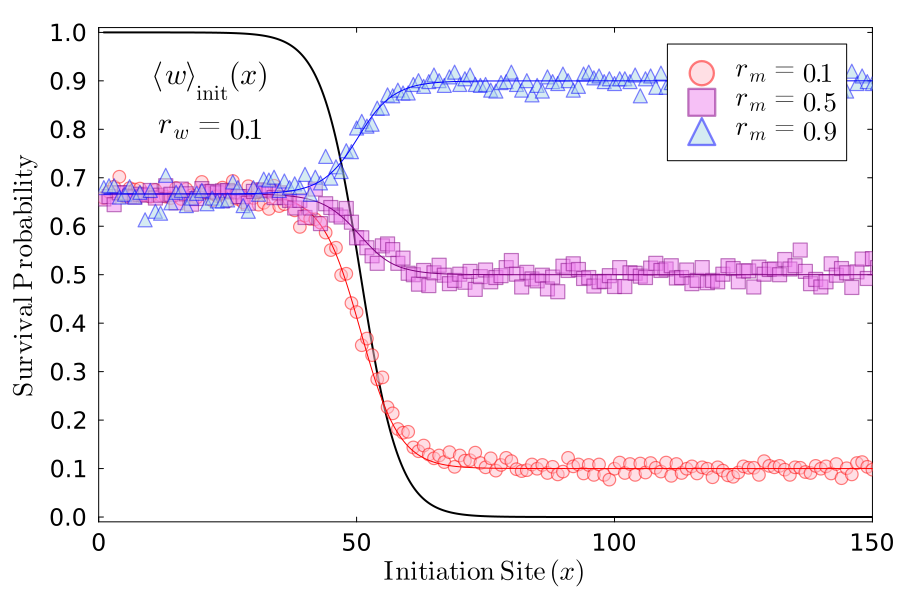

### paper_rw_0.1_Pwmm_-1.0_Pmwm_1.0_no_num_psurv_multiple_rms.png

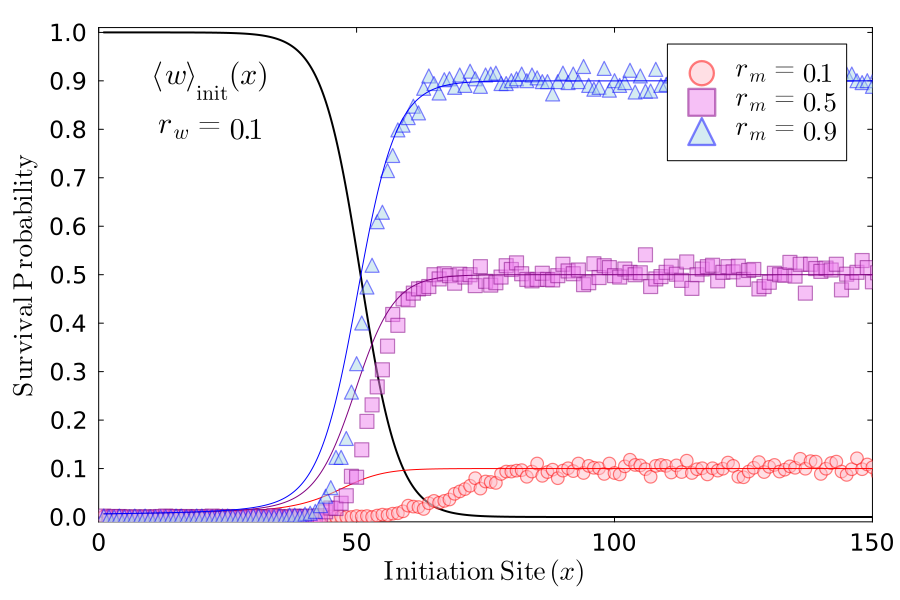

### paper_rw_0.5_Pwmm_0.5_Pmwm_0.75_no_num_psurv_multiple_rms.png

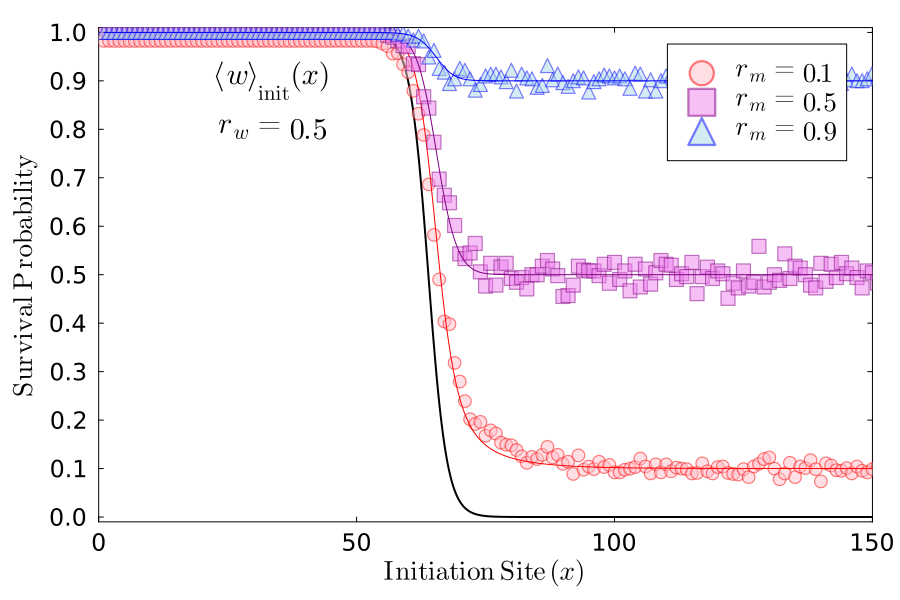

### paper_rw_0.5_Pwmm_0.5_Pmwm_-0.25_no_num_psurv_multiple_rms.png

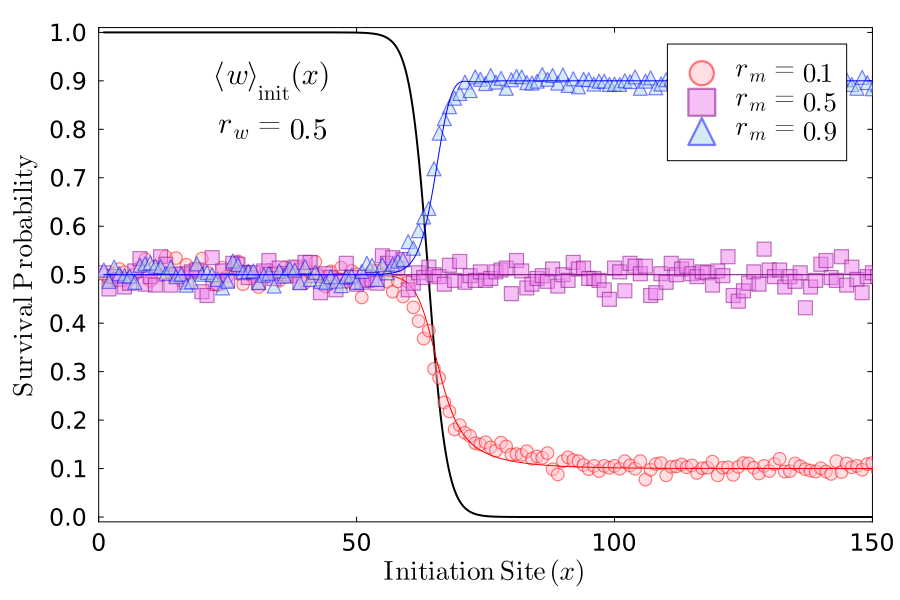

### paper_rw_0.5_Pwmm_-0.25_Pmwm_0.75_no_num_psurv_multiple_rms.png

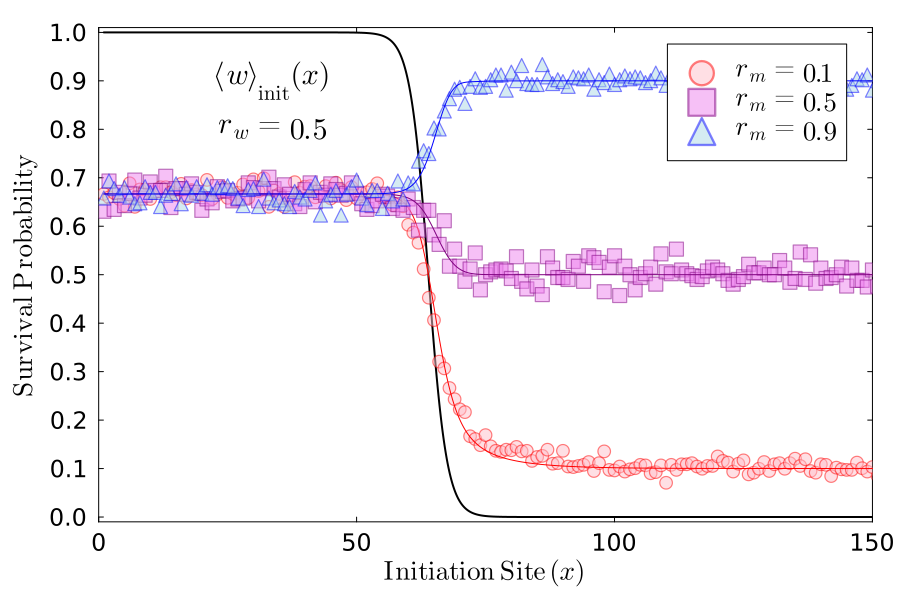

### paper_rw_0.5_Pwmm_-1.0_Pmwm_1.0_no_num_psurv_multiple_rms.png

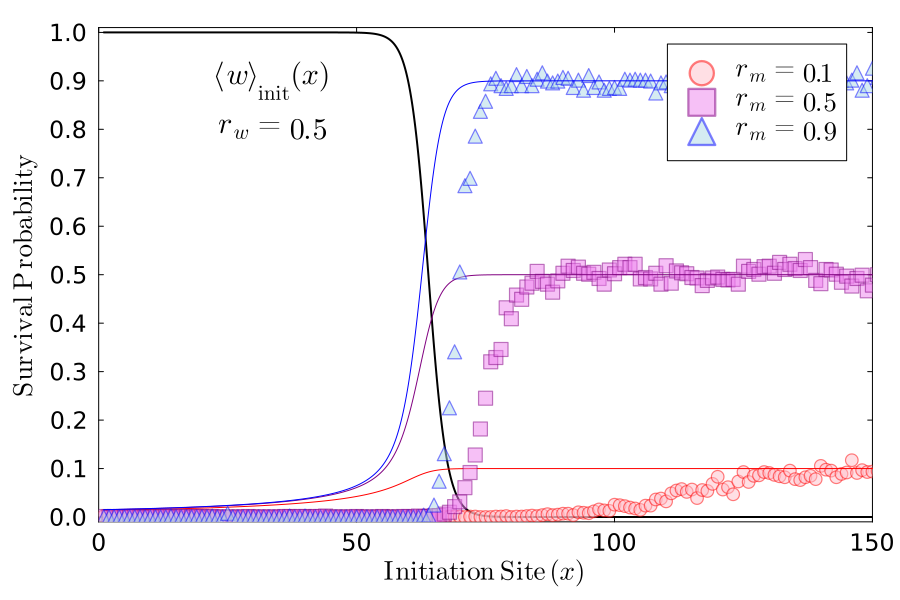

### paper_rw_0.9_Pwmm_0.5_Pmwm_0.75_no_num_psurv_multiple_rms.png

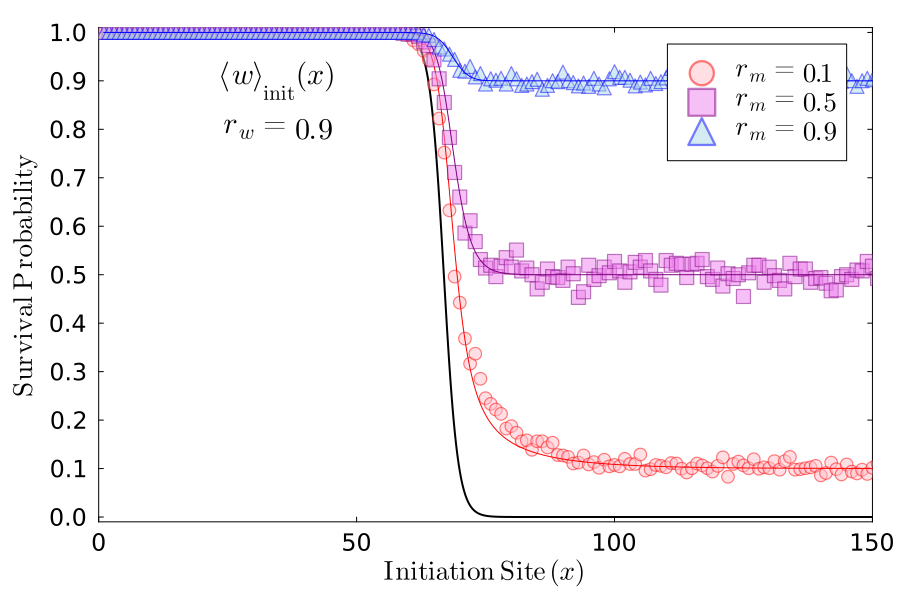

### paper_rw_0.9_Pwmm_0.5_Pmwm_-0.25_no_num_psurv_multiple_rms.png

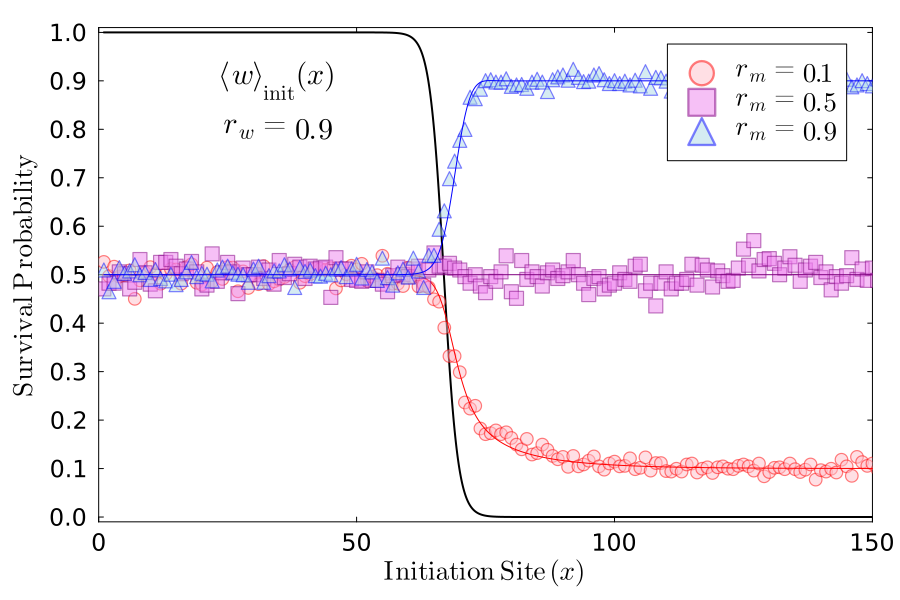

### paper_rw_0.9_Pwmm_-0.25_Pmwm_0.75_no_num_psurv_multiple_rms.png

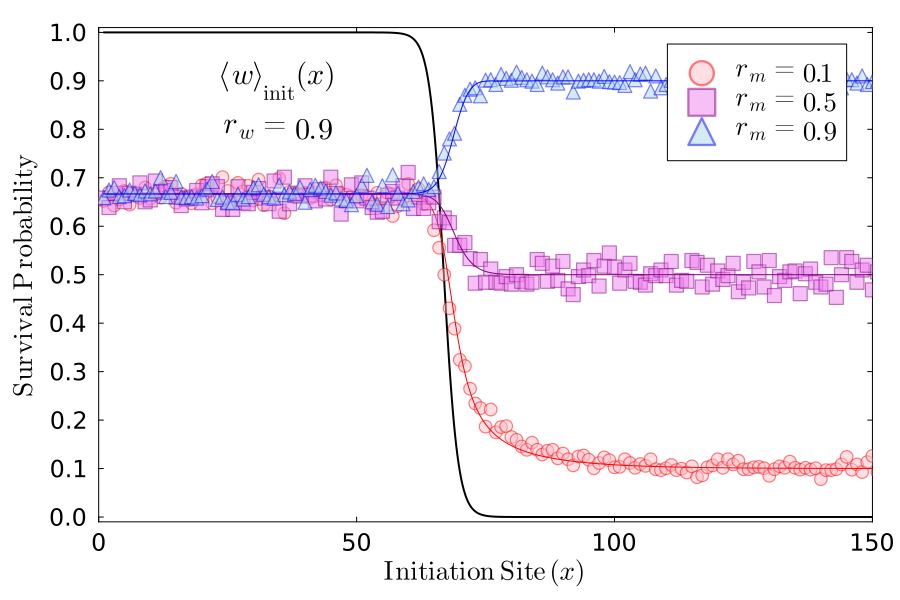

### paper_rw_0.9_Pwmm_-1.0_Pmwm_1.0_no_num_psurv_multiple_rms.png

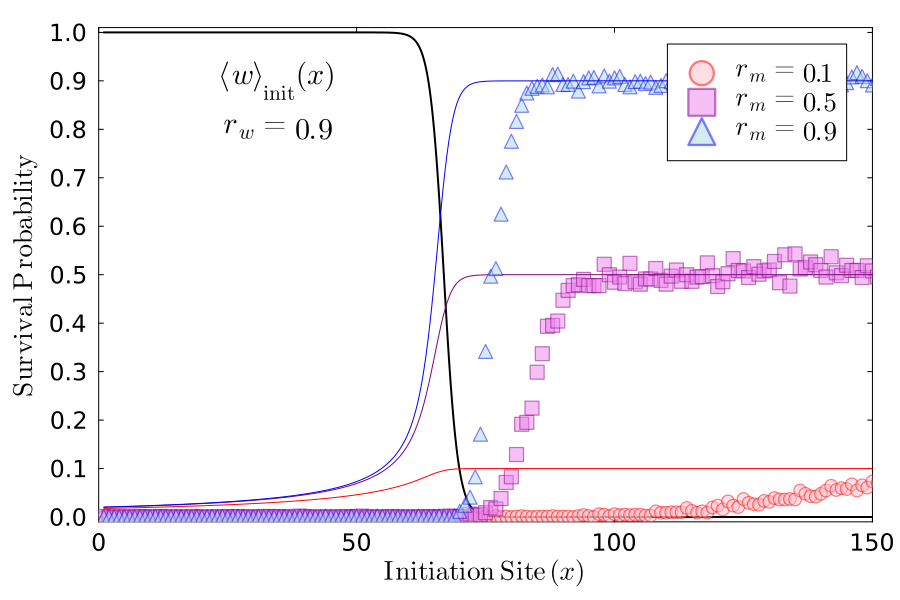

### Surf_vs_Abide_rw_0.1_rm_0.1_Pwmm_0.5_Pmwm_0.75.png

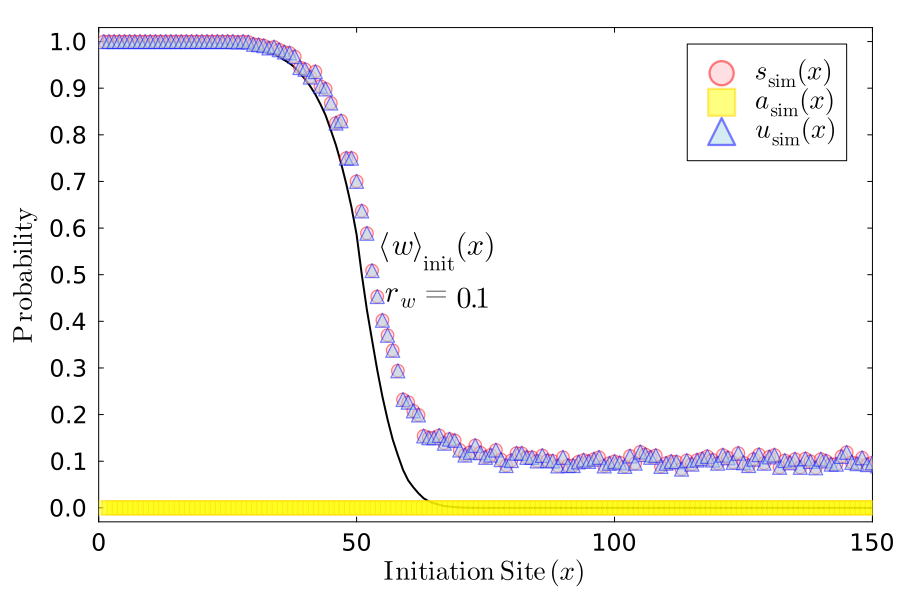

### Surf_vs_Abide_rw_0.1_rm_0.1_Pwmm_0.5_Pmwm_-0.25.png

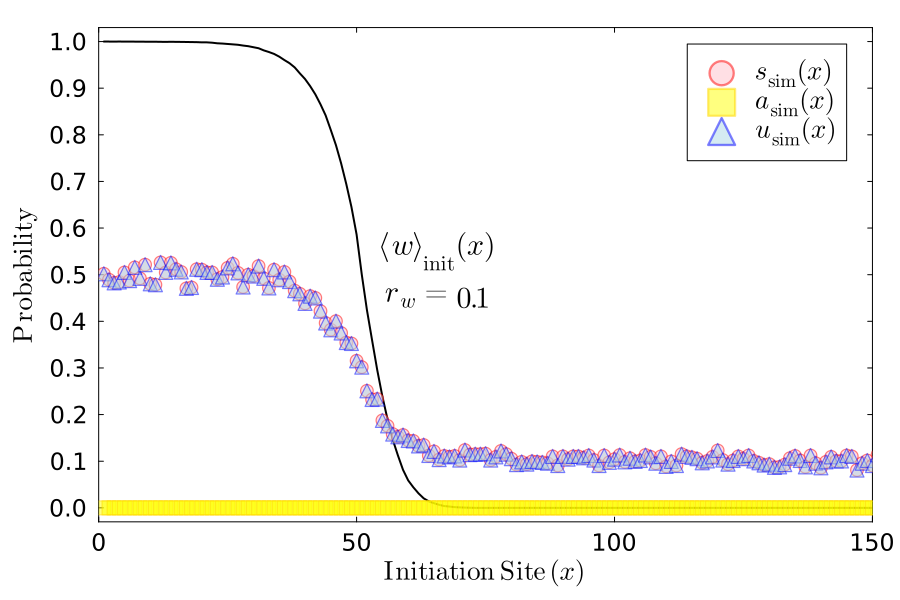

### Surf_vs_Abide_rw_0.1_rm_0.1_Pwmm_-0.25_Pmwm_0.75.png

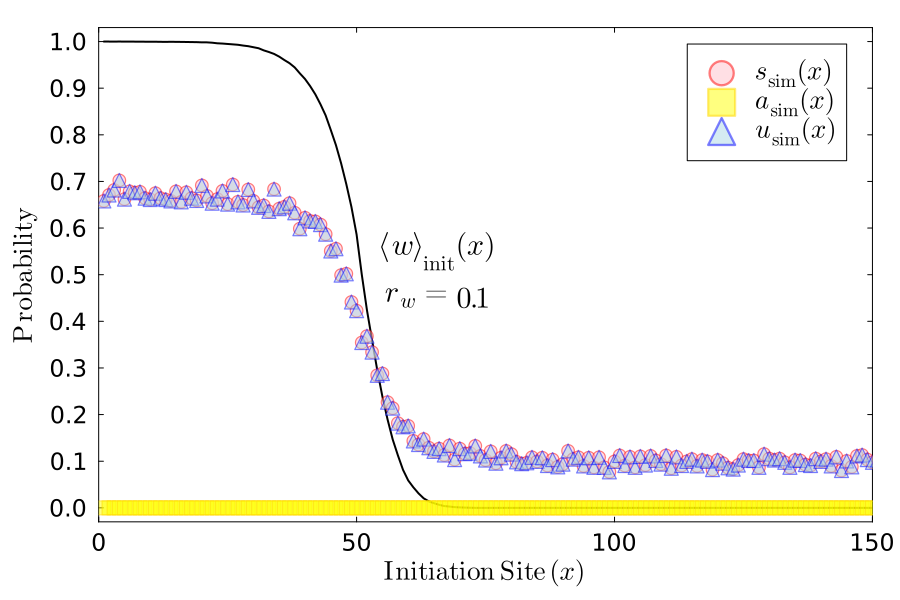

### Surf_vs_Abide_rw_0.1_rm_0.1_Pwmm_-1.0_Pmwm_1.0.png

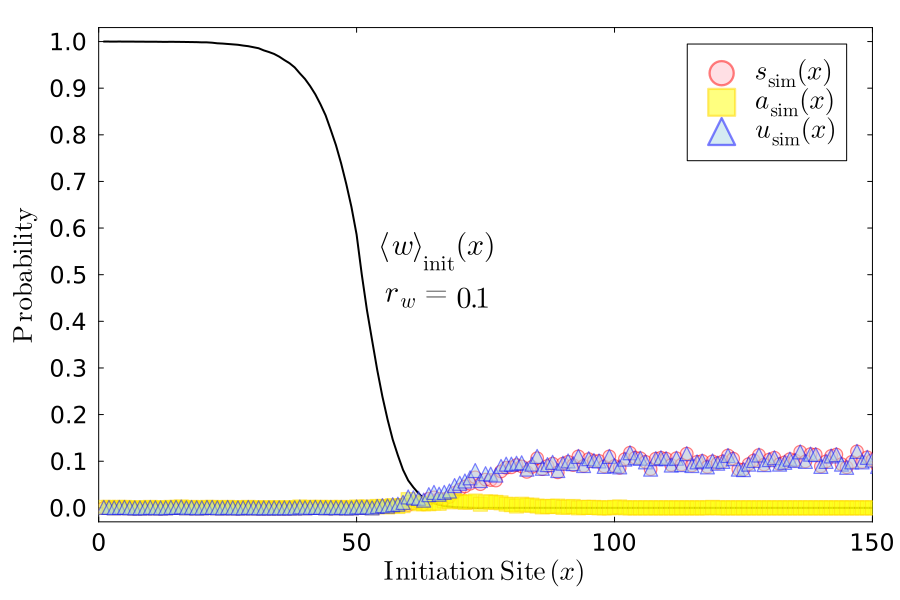

### Surf_vs_Abide_rw_0.1_rm_0.5_Pwmm_0.5_Pmwm_0.75.png

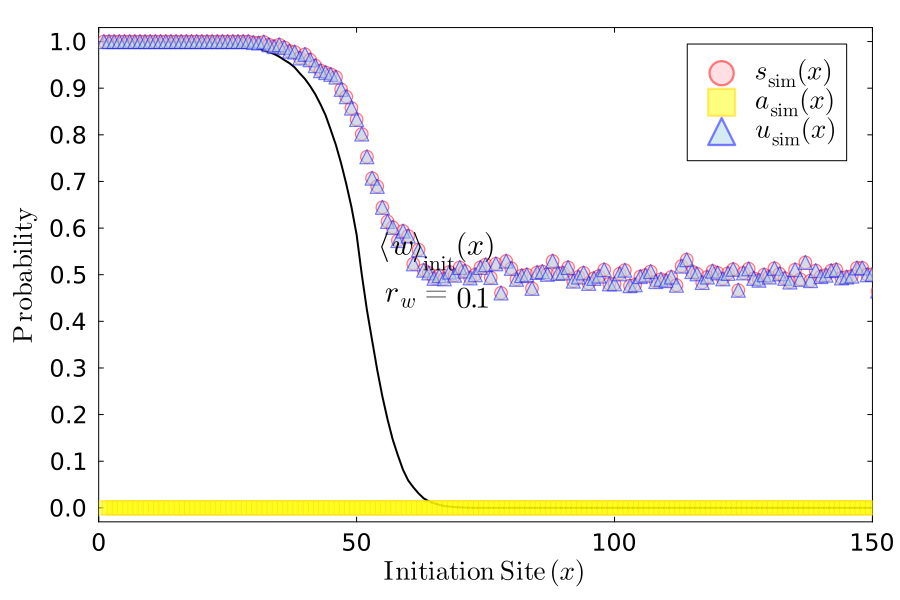

### Surf_vs_Abide_rw_0.1_rm_0.5_Pwmm_0.5_Pmwm_-0.25.png

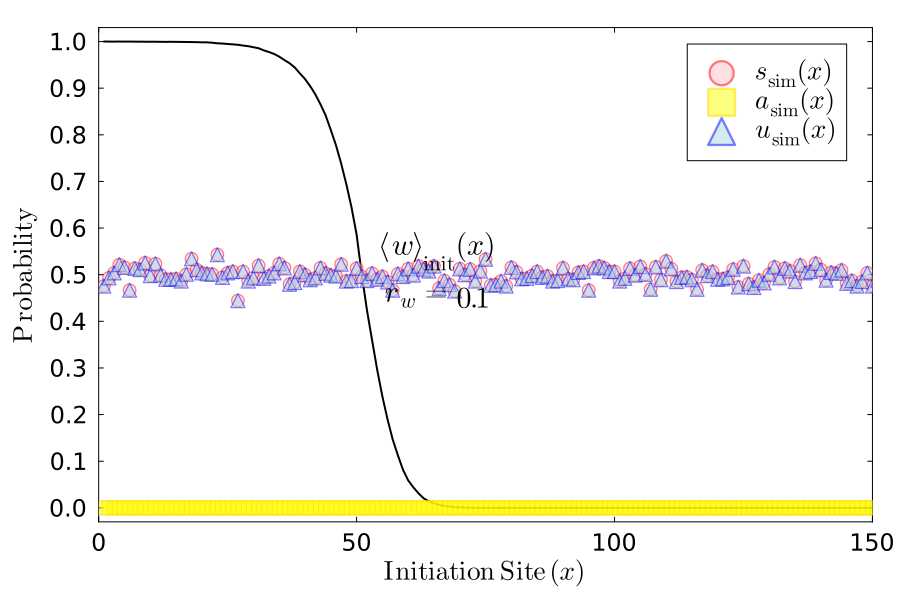

### Surf_vs_Abide_rw_0.1_rm_0.5_Pwmm_-0.25_Pmwm_0.75.png

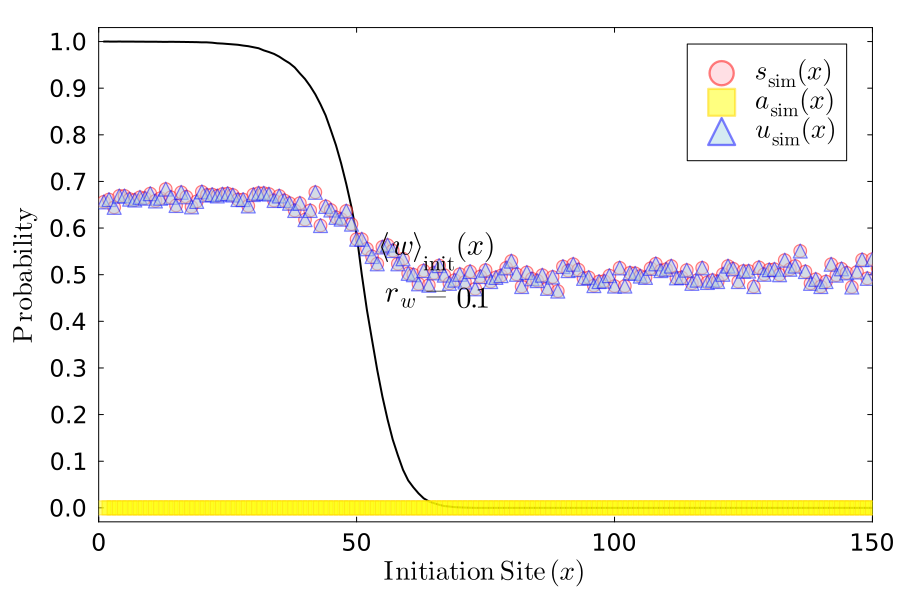

### Surf_vs_Abide_rw_0.1_rm_0.5_Pwmm_-1.0_Pmwm_1.0.png

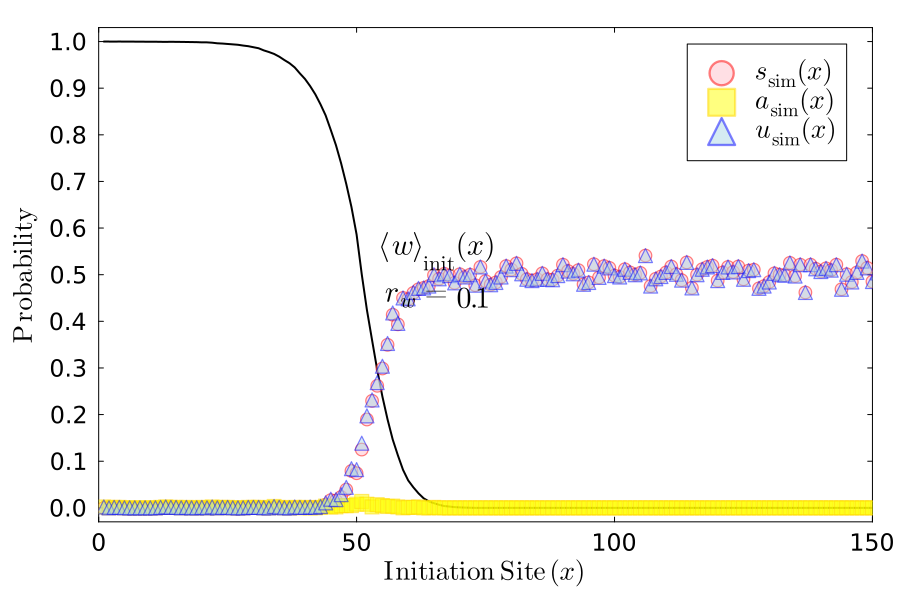

### Surf_vs_Abide_rw_0.1_rm_0.9_Pwmm_0.5_Pmwm_0.75.png

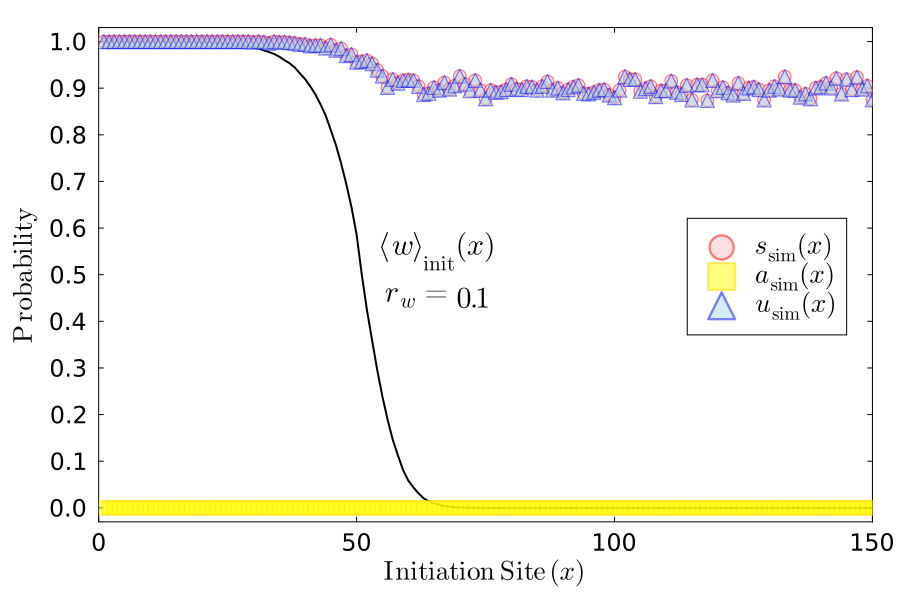

### Surf_vs_Abide_rw_0.1_rm_0.9_Pwmm_0.5_Pmwm_-0.25.png

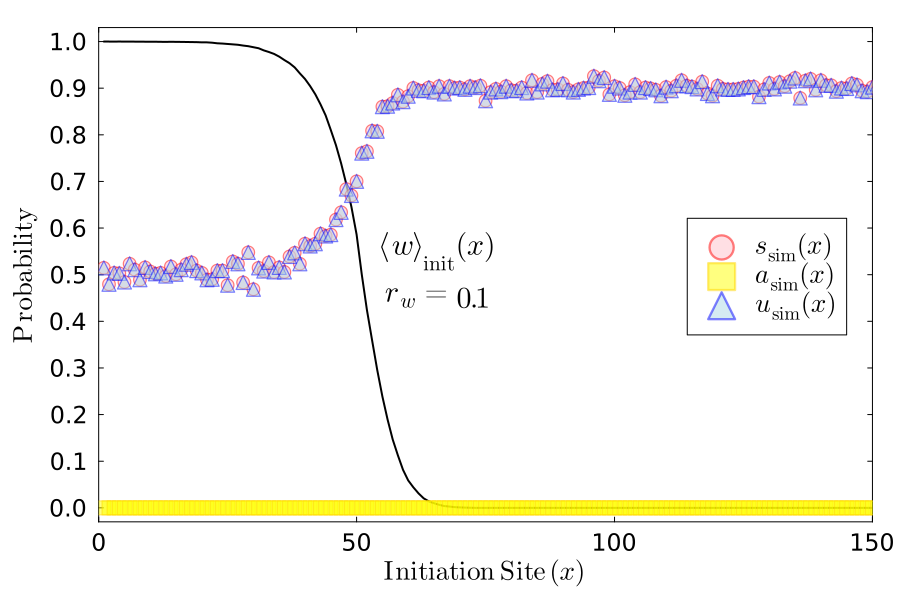

### Surf_vs_Abide_rw_0.1_rm_0.9_Pwmm_-0.25_Pmwm_0.75.png

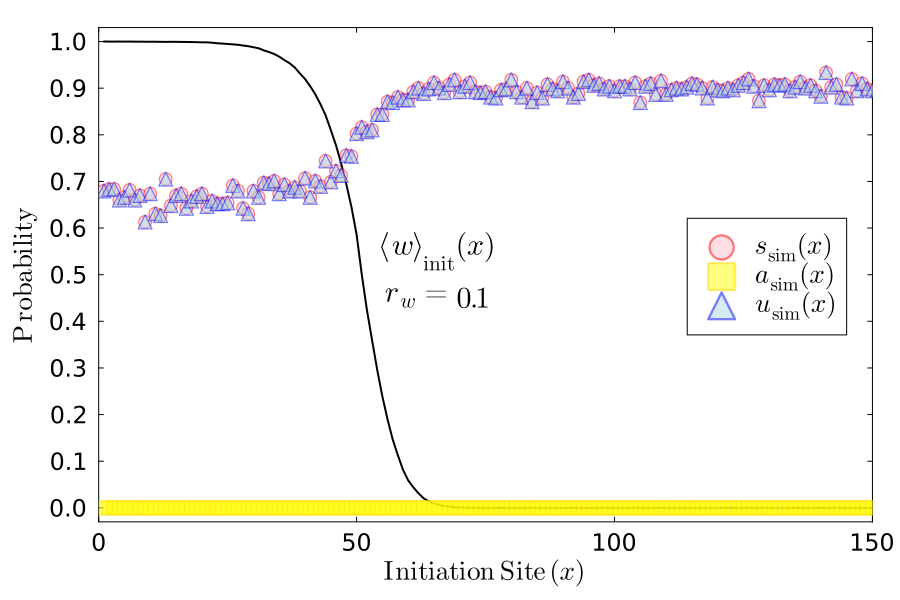

### Surf_vs_Abide_rw_0.1_rm_0.9_Pwmm_-1.0_Pmwm_1.0.png

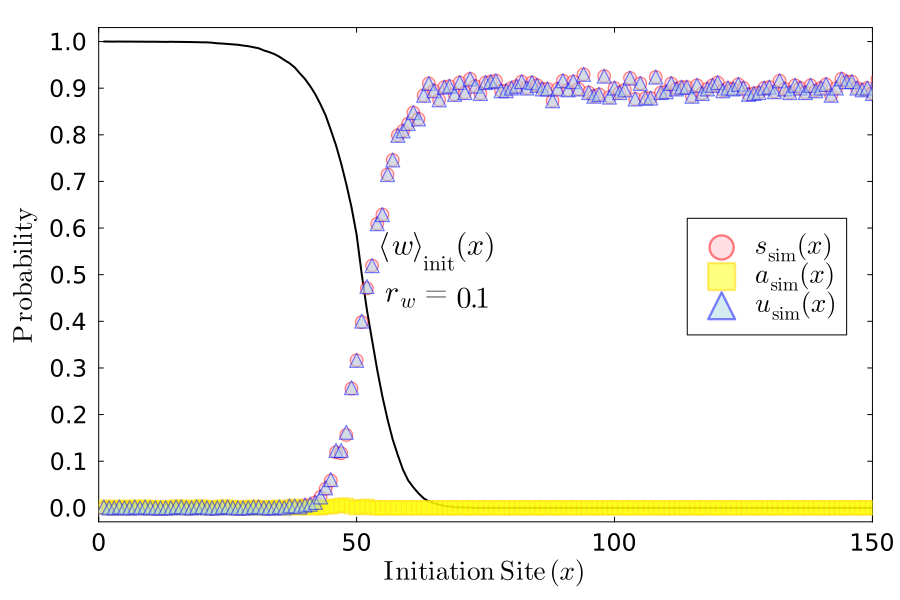

### Surf_vs_Abide_rw_0.5_rm_0.1_Pwmm_0.5_Pmwm_0.75.png

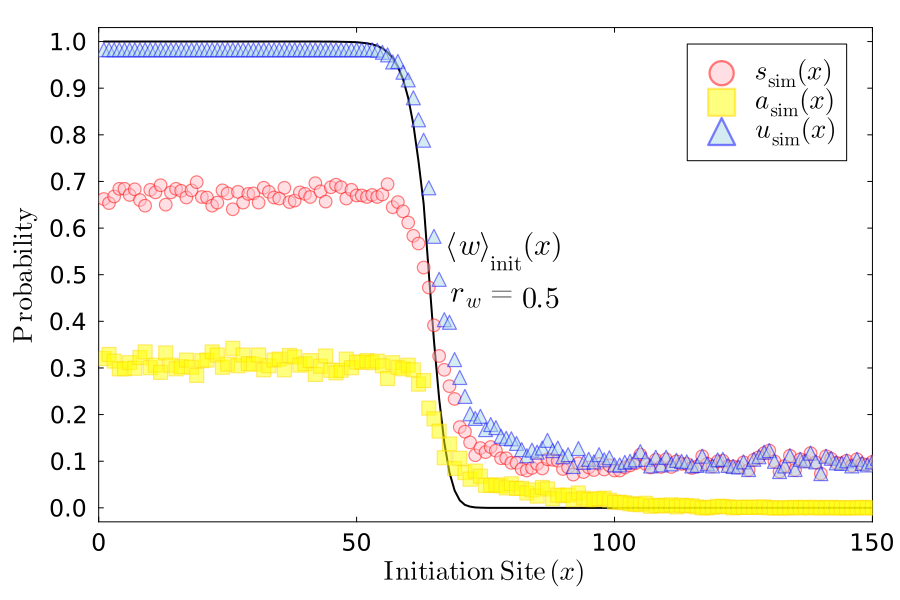

### Surf_vs_Abide_rw_0.5_rm_0.1_Pwmm_0.5_Pmwm_-0.25.png

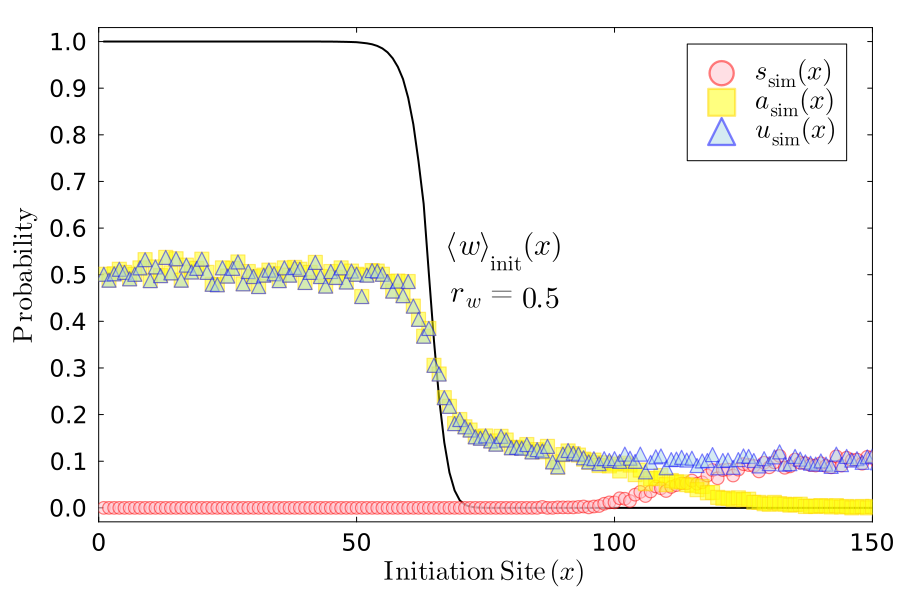

### Surf_vs_Abide_rw_0.5_rm_0.1_Pwmm_-0.25_Pmwm_0.75.png

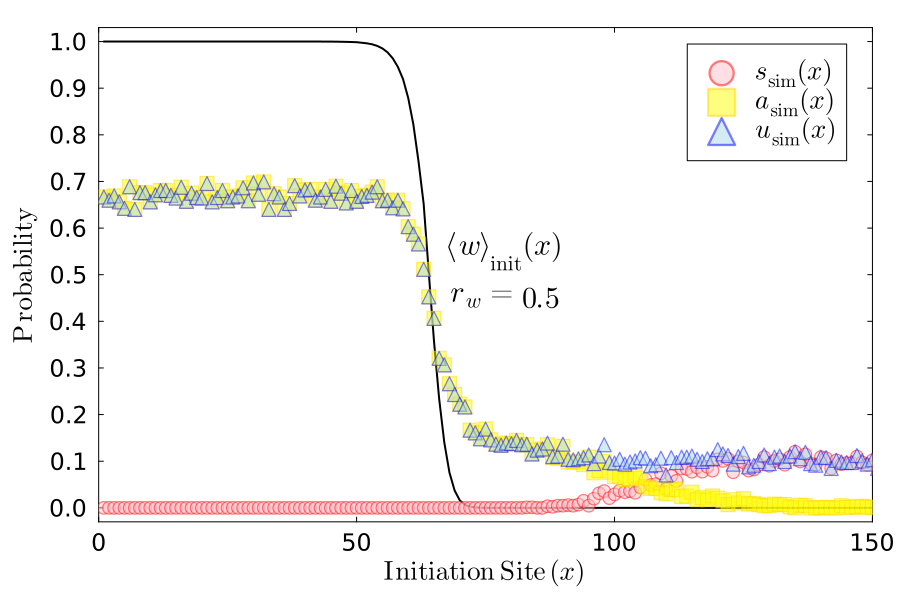

### Surf_vs_Abide_rw_0.5_rm_0.1_Pwmm_-1.0_Pmwm_1.0.png

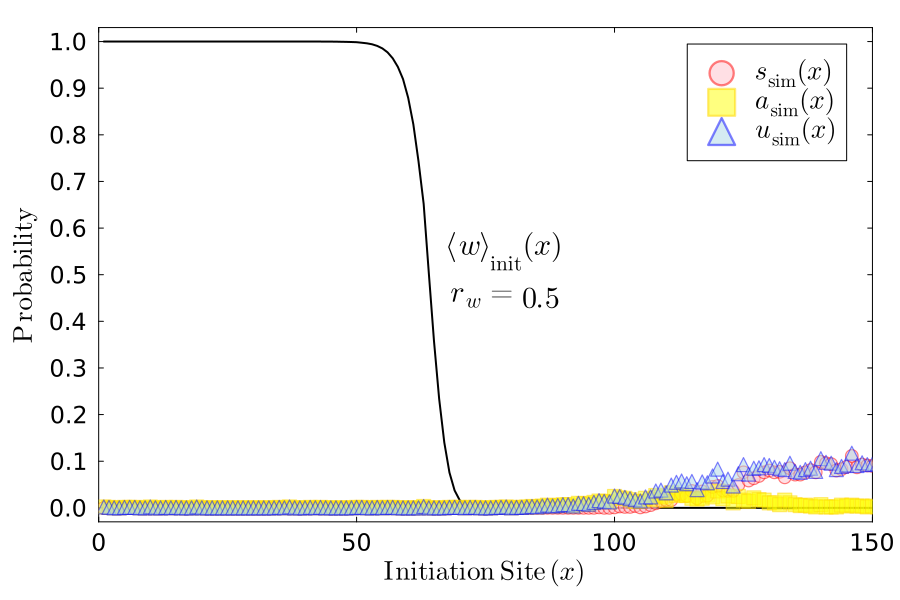

### Surf_vs_Abide_rw_0.5_rm_0.5_Pwmm_0.5_Pmwm_0.75.png

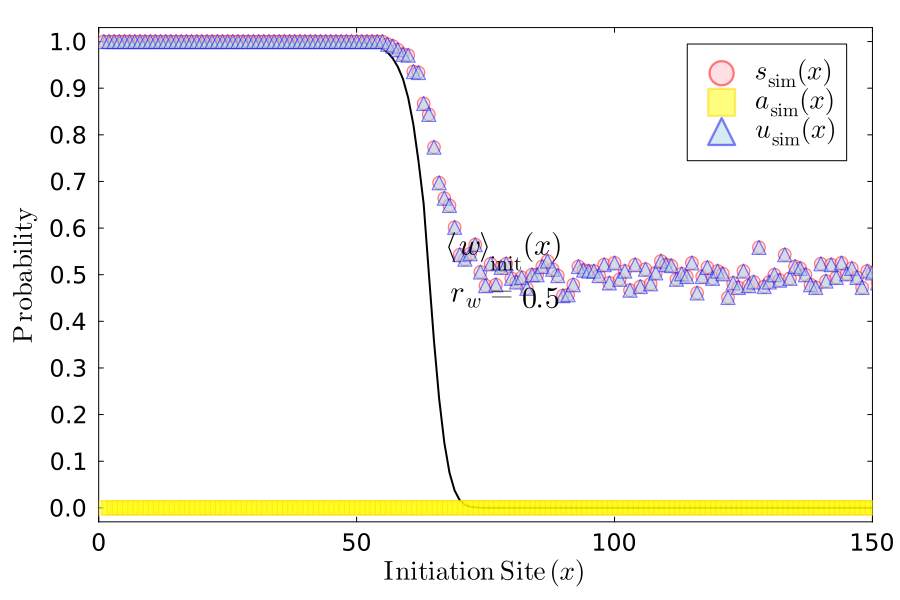

### Surf_vs_Abide_rw_0.5_rm_0.5_Pwmm_0.5_Pmwm_-0.25.png

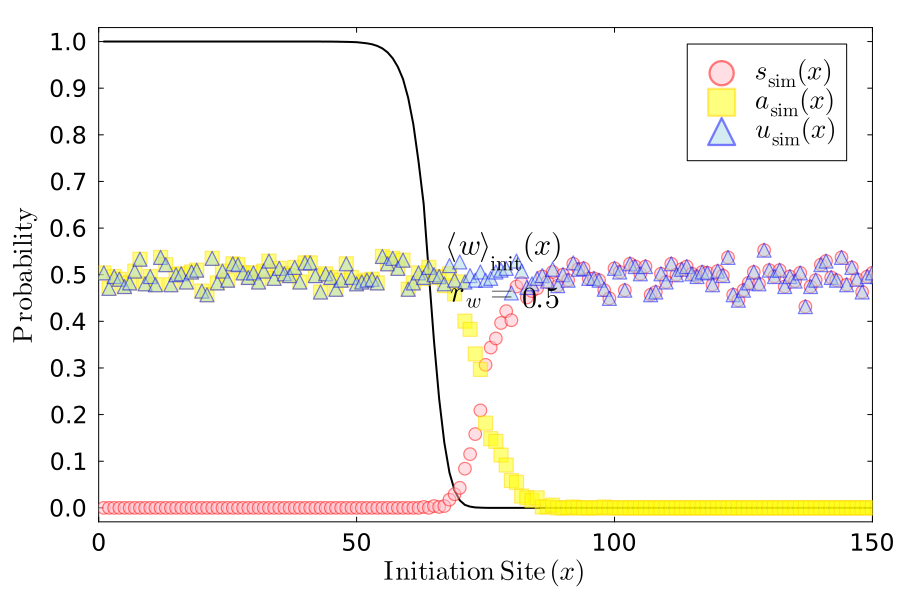

### Surf_vs_Abide_rw_0.5_rm_0.5_Pwmm_-0.25_Pmwm_0.75.png

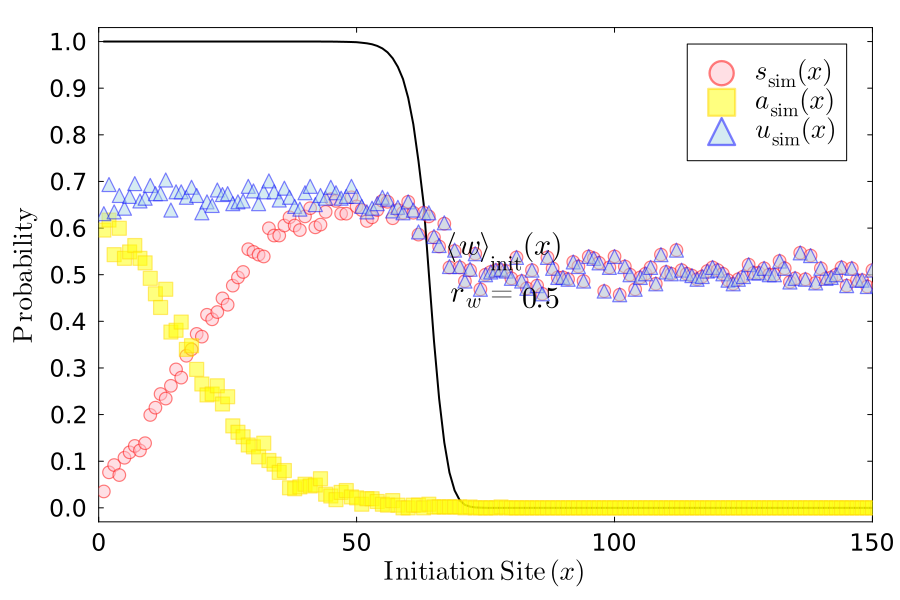

### Surf_vs_Abide_rw_0.5_rm_0.5_Pwmm_-1.0_Pmwm_1.0.png

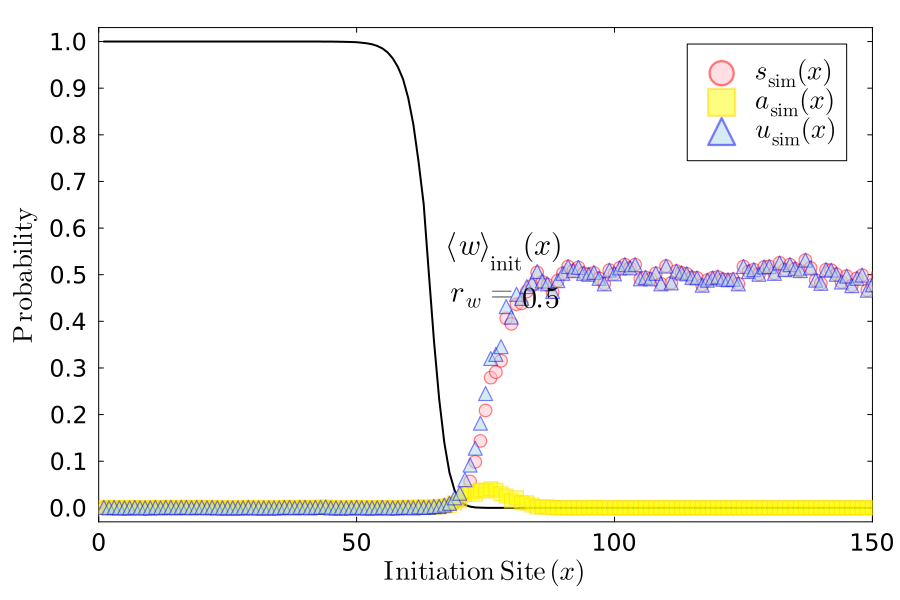

### Surf_vs_Abide_rw_0.5_rm_0.9_Pwmm_0.5_Pmwm_0.75.png

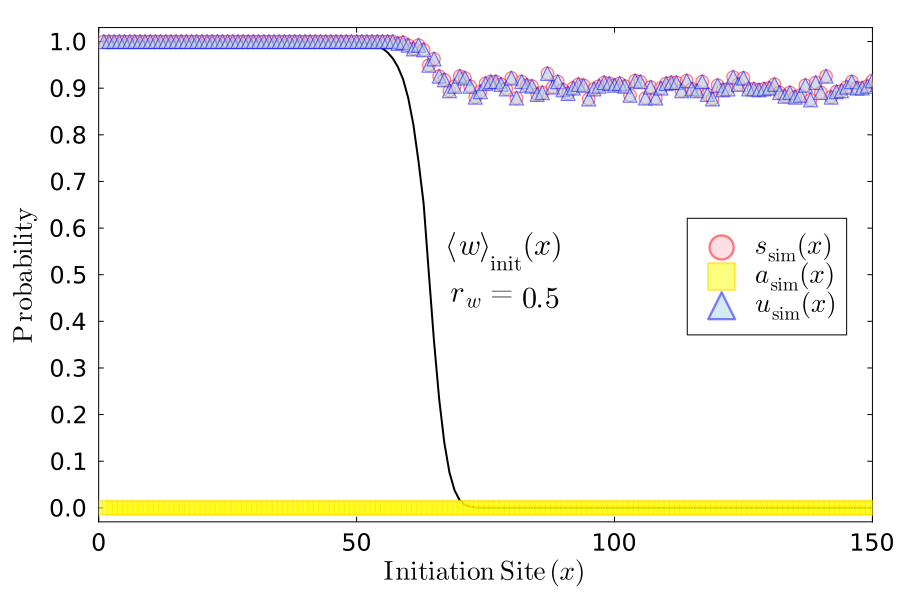

### Surf_vs_Abide_rw_0.5_rm_0.9_Pwmm_0.5_Pmwm_-0.25.png

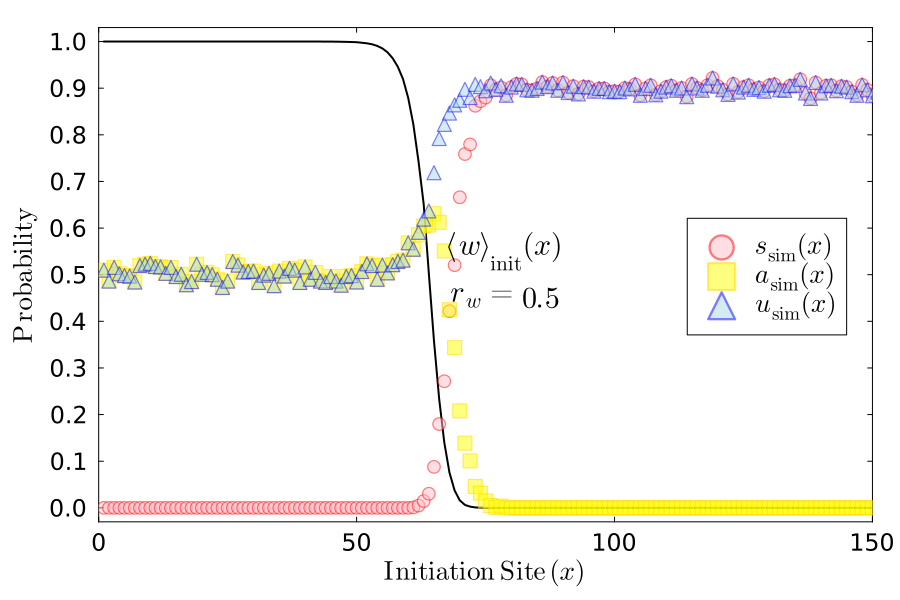

### Surf_vs_Abide_rw_0.5_rm_0.9_Pwmm_-0.25_Pmwm_0.75.png

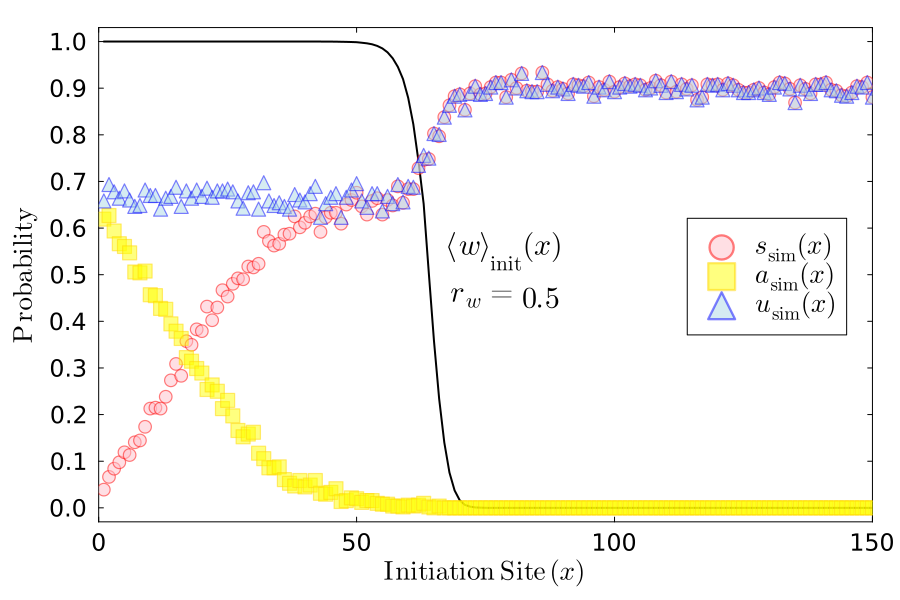

### Surf_vs_Abide_rw_0.5_rm_0.9_Pwmm_-1.0_Pmwm_1.0.png

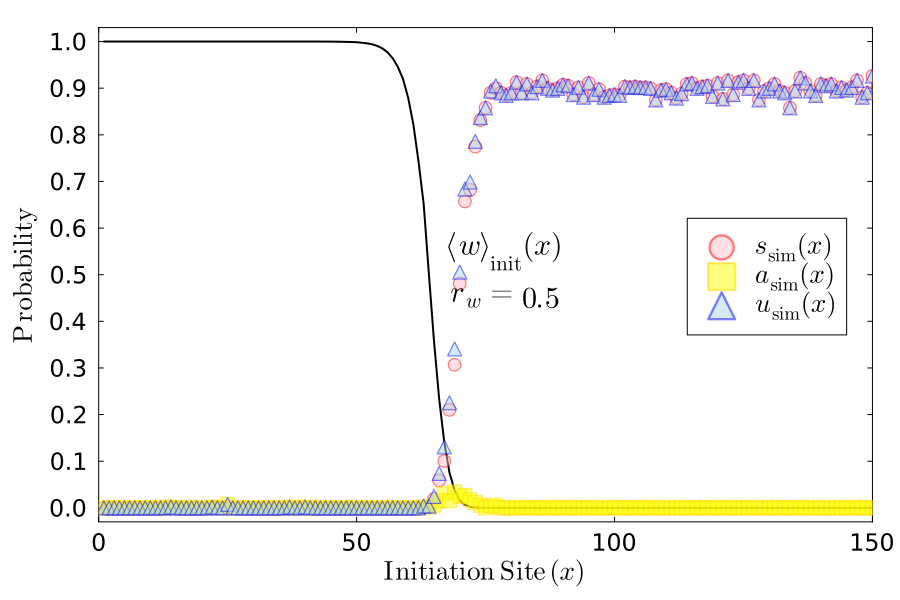

### Surf_vs_Abide_rw_0.9_rm_0.1_Pwmm_0.5_Pmwm_0.75.png

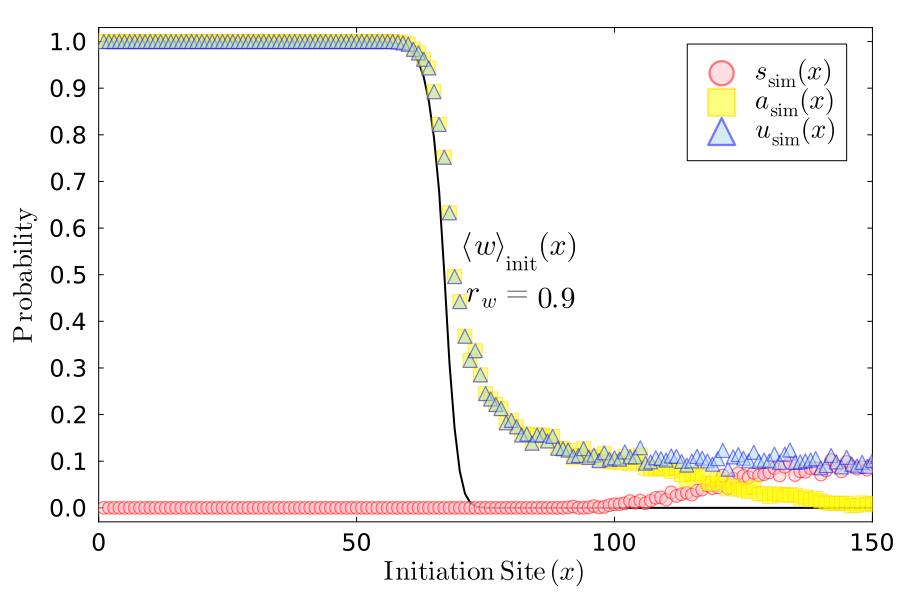

### Surf_vs_Abide_rw_0.9_rm_0.1_Pwmm_0.5_Pmwm_-0.25.png

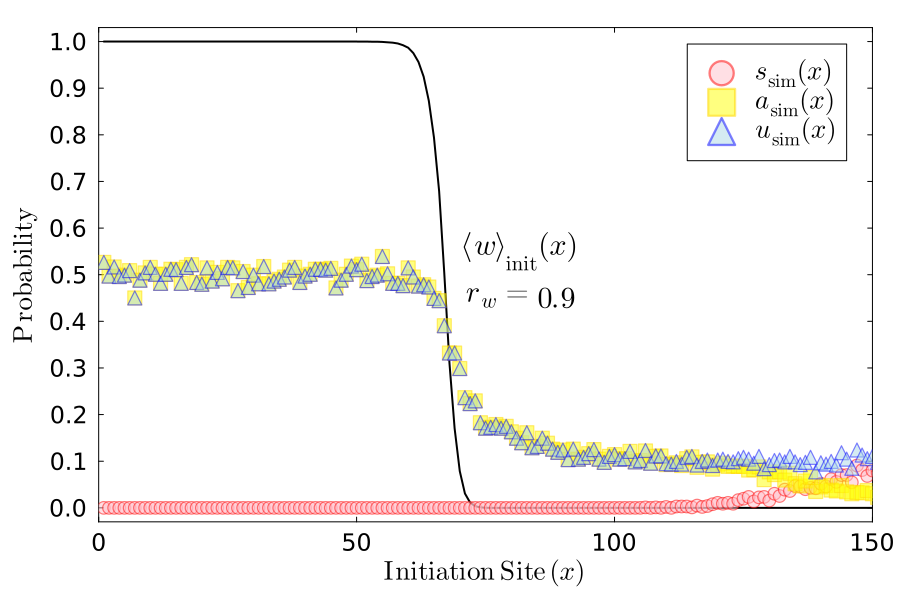

### Surf_vs_Abide_rw_0.9_rm_0.1_Pwmm_-0.25_Pmwm_0.75.png

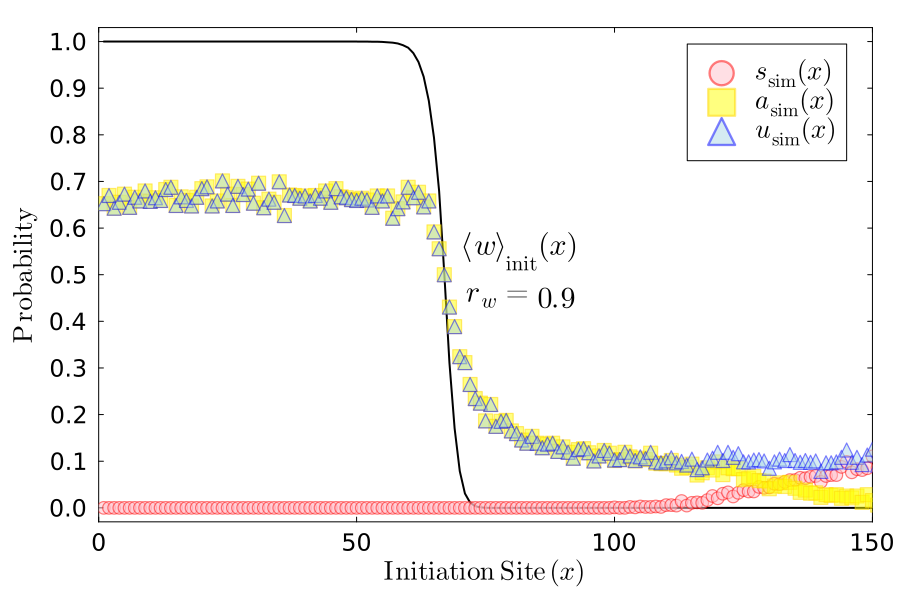

### Surf_vs_Abide_rw_0.9_rm_0.1_Pwmm_-1.0_Pmwm_1.0.png

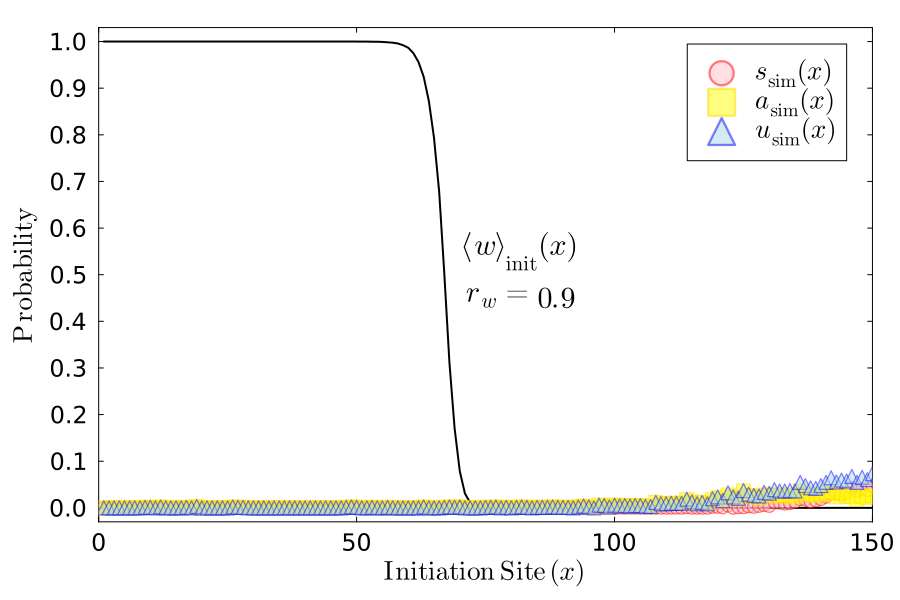

### Surf_vs_Abide_rw_0.9_rm_0.5_Pwmm_0.5_Pmwm_0.75.png

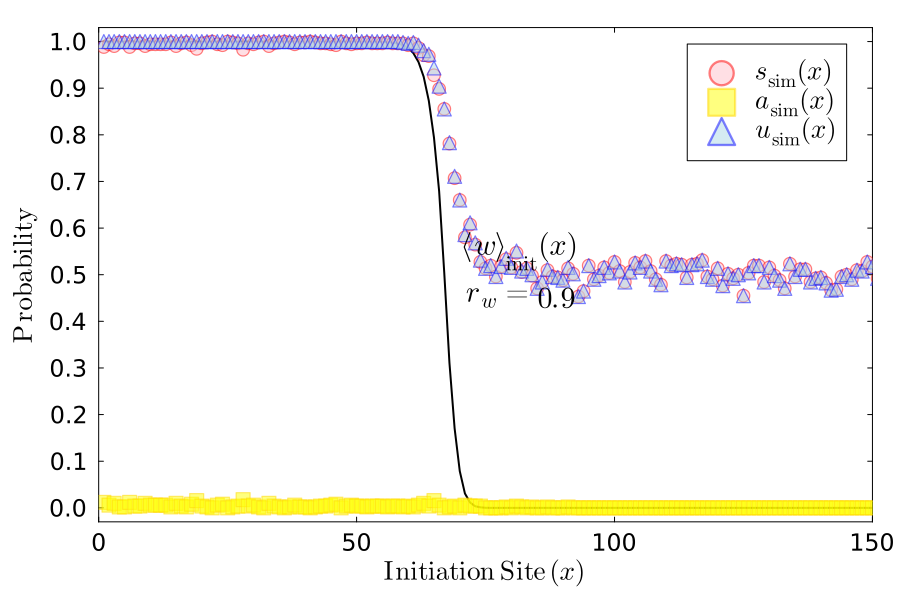

### Surf_vs_Abide_rw_0.9_rm_0.5_Pwmm_0.5_Pmwm_-0.25.png

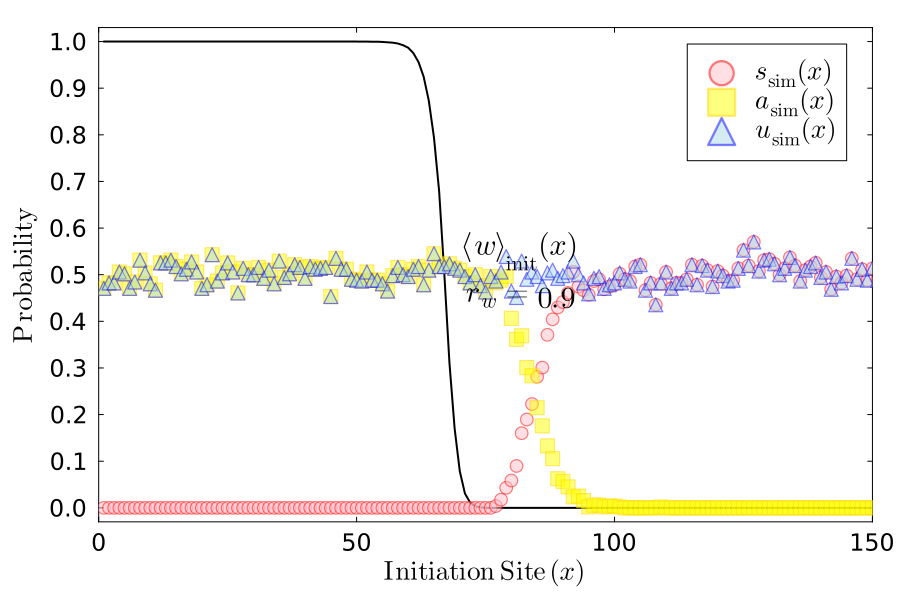

### Surf_vs_Abide_rw_0.9_rm_0.5_Pwmm_-0.25_Pmwm_0.75.png

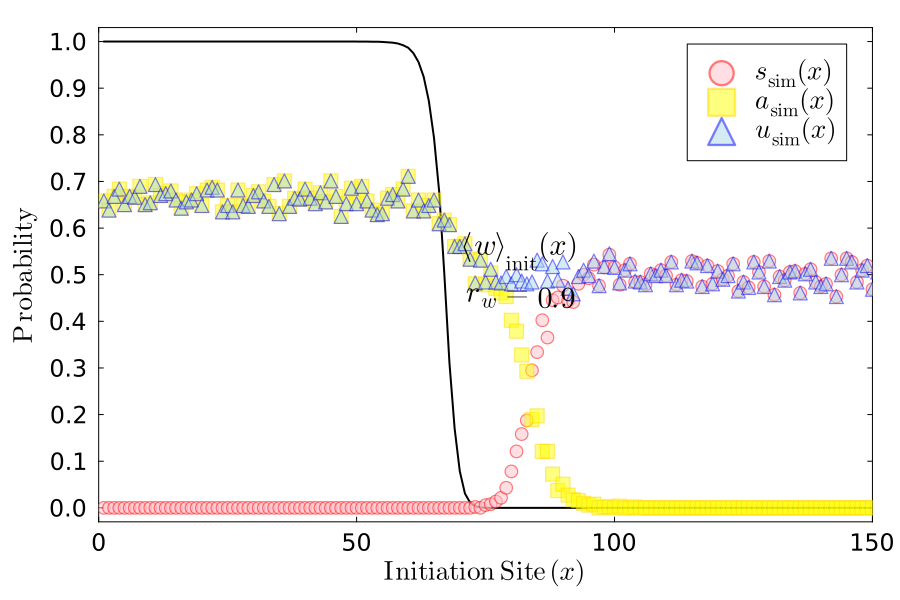

### Surf_vs_Abide_rw_0.9_rm_0.5_Pwmm_-1.0_Pmwm_1.0.png

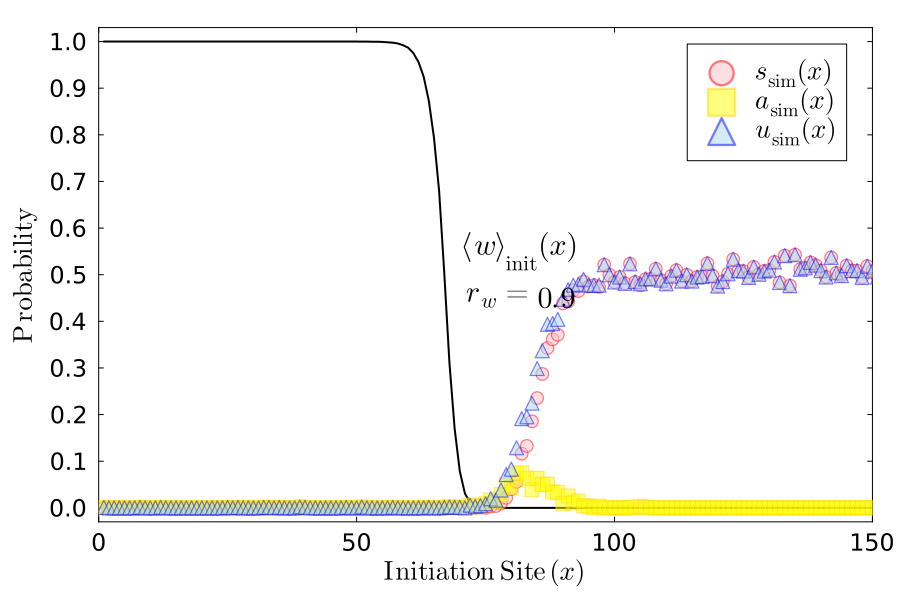

### Surf_vs_Abide_rw_0.9_rm_0.9_Pwmm_0.5_Pmwm_0.75.png

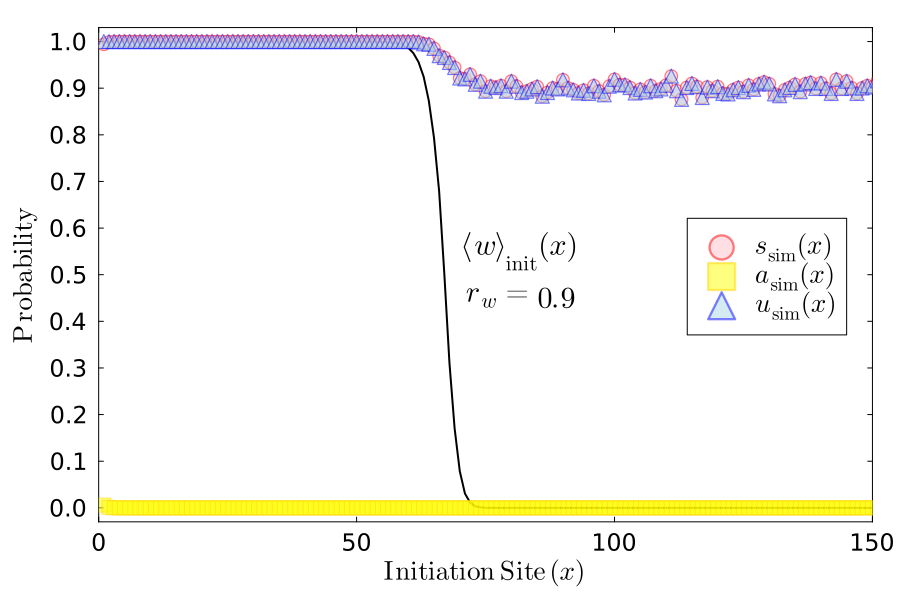

### Surf_vs_Abide_rw_0.9_rm_0.9_Pwmm_0.5_Pmwm_-0.25.png

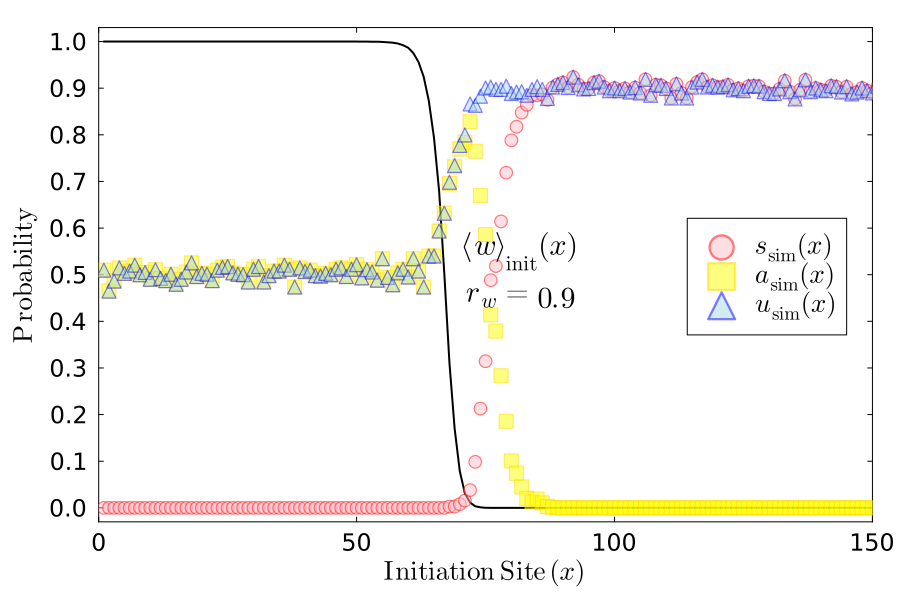

### Surf_vs_Abide_rw_0.9_rm_0.9_Pwmm_-0.25_Pmwm_0.75.png

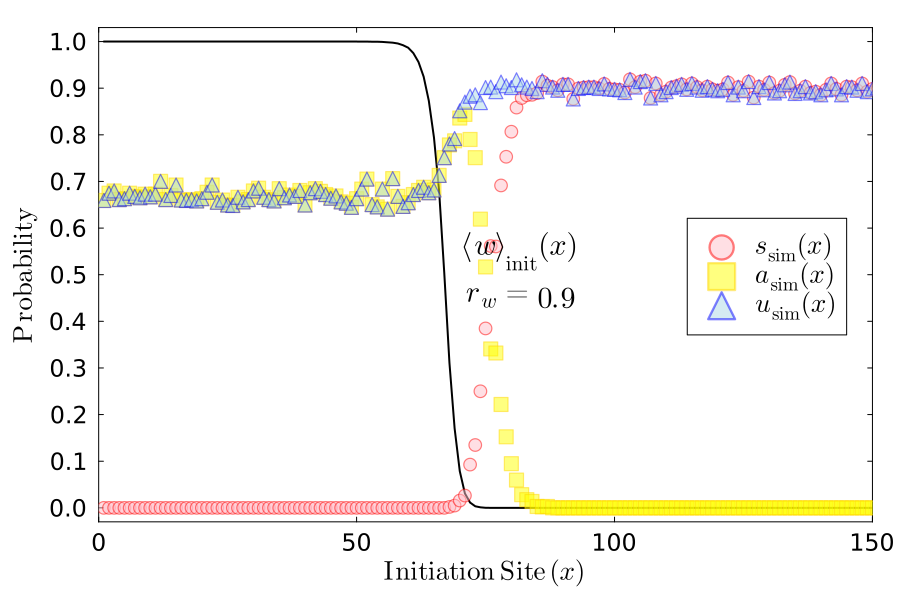

### Surf_vs_Abide_rw_0.9_rm_0.9_Pwmm_-1.0_Pmwm_1.0.png

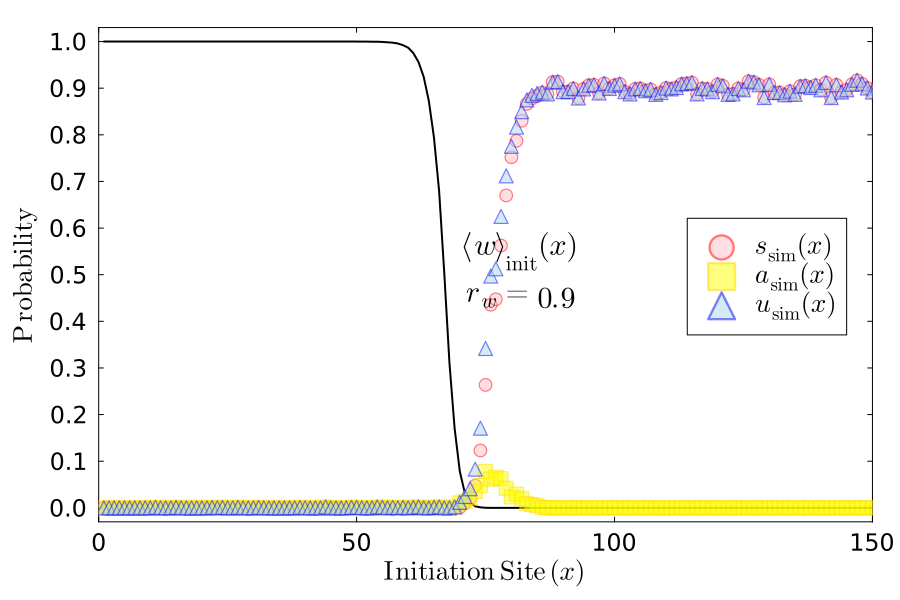
